# Using Progressive Context Encoders for Anomaly Detection in Digital Pathology Images

**DOI:** 10.1101/2021.07.02.450957

**Authors:** Ryan Gillard, Chady Meroueh, Qiangqiang Gu, Naresh Prodduturi, Sandhya Patil, Thomas J Flotte, Steven N Hart

**Affiliations:** Google, Mountain View, CA; Mayo Clinic, Rochester, MN; Mayo Clinic Rochester, MN

## Abstract

Whole slide imaging (WSI) is transforming the practice of pathology, converting a qualitative discipline into a quantitative one. However, one must exercise caution in interpreting algorithm assertions, particularly in pathology where an incorrect classification could have profound impacts on a patient, and rare classes exist that may not have been seen by the algorithm during training. A more robust approach would be to identify areas of an image for which the pathologist should concentrate their effort to make a final diagnosis. This anomaly detection strategy would be ideal for WSI, but given the extremely high resolution and large file sizes, such an approach is difficult. Here, we combine progressive generative adversarial networks with a flexible adversarial autoencoder architecture capable of learning the “normal distribution” of WSIs of normal skin tissue at extremely high resolution and demonstrate its anomaly detection performance. Our approach yielded pixel-level accuracy of 89% for identifying melanoma, suggesting that our label-free anomaly detection pipeline is a viable strategy for generating high quality annotations - without tedious manual segmentation by pathologists. The code is publicly available at https://github.com/Steven-N-Hart/P-CEAD.

## MAIN

Skin cancer is the most common of all human cancers, with 1 million people in the United States diagnosed each year with some type of the disease. Most skin cancers are basal and squamous cell carcinomas. While malignant, these types are relatively easily cured with minimal surgical intervention. Malignant melanomas, however, account for about 1% of all skin cancers in the United States, but cause the majority of skin cancer deaths. The number of people diagnosed with melanoma has risen sharply over the past 3 decades. In men and women ages 50 and older, the number of people diagnosed with melanoma increased 3% per year from 2006 to 2015. Identification and validation of melanomas are of critical importance, as patients with dermatologist-detected melanomas have better survival, lower overall mortality, and lower cancer-related mortality^1^.

The most important recent advances in microscopy for surgical pathology were the invention of the digital microscope^2^ and whole slide imaging (WSI)^3^. The current generation of high-speed, high-capacity whole slide scanners can process between 1 and 1,000 slides at multiple resolutions and different image planes (i.e. z-stacks). The quality of the images has been steadily increasing over time. Several recent comparisons have been made between rendering a diagnosis on a glass slide and a digital assessment, with concordances reported between 75-100%^4–6^. The image files themselves are quite large, between 2-5 GB each, requiring analysis to be conducted on much smaller regions (a.k.a. “patches”)^7^.

Perhaps the most exciting opportunity resulting from the digital pathology transition to WSI is the potential utility of applying artificial intelligence (AI) allowing for computational pathology^8–10^. We previously showed that AI was capable of differentiating between Spitz and Conventional Nevi (benign skin lesions)^11^. Later, Hekler *et al*.,^12^ trained an AI to differentiate between compound or junctional nevi and melanoma. However, both approaches suffer from limitations of the training set. The World Health Organization recognizes 9 different types of melanoma^13^, but there are also significant numbers of benign lesions that mimic melanomas or are detected at premalignant stages^14^. Models like these that only account for two possible outcomes (Spitz/Conventional or benign/malignant), necessarily means that if the input image is not from either class, then the model will always make an incorrect diagnosis. Given some of the more rare examples of benign and malignant lesions, it may not be possible to accumulate enough examples from each class to build a model that captures everything one would see in dermatopathology practice.

A more practical approach would be to convert the “classification” problem into an “anomaly detection” problem. The major difference is that during training, the anomaly detection approach only sees one class (e.g. normal) whereas the classification approach needs examples from each possible class. In digital pathology, the former is ideal since examples with negative findings are easily acquired. Rather than learning how to differentiate between all possible classes, the model instead learns to weigh each pixel as to how likely it belongs to the normal class, and if that score lies outside some expected distribution, then it gets flagged as an anomaly - without the need to say what class that anomaly belongs to. The advantage for digital pathology is that the areas of interest can quickly be identified and carefully scrutinized by the pathologist - who can more carefully consider the question of “what” the anomaly actually is.

### Anomaly Detection using GANs

Generative adversarial networks (GANs) are well suited for anomaly detection problems. A GAN consists of two adversarial modules, a generator (*G*) and a discriminator (*D*). G typically learns to generate realistic looking images 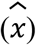 from latent-space vectors (*z*), which are then served to the discriminator for determining their real or fake status (*q*). However the mapping from 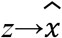 is different than 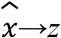 which is important for understanding how and where the vector contributes to the generated image. AnnoGAN^15^ requires an additional step to learn this mapping. To avoid this, EBGAN^16^ simultaneously added an encoder (*E*) model for joint training of 3 networks (*G, D, E*) and added the latent/pseudo-latent variable (*z*/*z’*) as an input into *D* as in BiGAN^17^. GANomaly^18^ rearranged the models into an adversarial autoencoder wherein 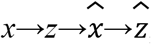, thereby removing the true *G* and leaving a bowtie architecture generator that both encodes (*G_E_*) and decodes (*G_D_*)). An adversarial loss 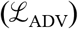 is calculated from a D, a contextual (a.k.a. reconstruction) loss 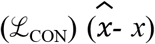, and an encoder loss 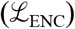 measuring the difference of latent space mappings 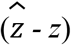. Di Mattia et al.,^19^ recently reviewed the use of each of these GANs for use in anomaly detection. The limitation to each of these methods is that they have been applied only to low resolution images. Berg *et al*.,^20^ proposed GANanomalyDetection (by combining ProGAN^21^ with a more updated version of AnnoGAN^15^), which theoretically could work for high resolution images, but was only tested on low resolution images. The main novelty of the ProGAN approach was the concept of training the model on smaller representations of the source images, while gradually increasing the image size and model architecture to very high resolution (1024×1024 pixels). The question as to whether the integration of progression learning in GANanomalyDetection for ultra high resolution images in digital pathology remains unanswered.

The work in GANomaly^18^ showed that one could ignore the complication of GANs altogether and use autoencoders (AEs). Autoencoders compress an image *x* to a small latent space *z* which is then transformed into a reconstructed image *x’*, while minimizing the reconstruction error between *x* and 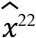^22^. Naturally, this ideal property led to the use of AEs for image compression^23^. Adversarial autoencoders however, take this one step further by drawing samples from the latent distribution *z* and combining the original reconstruction loss 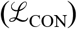 with an adversarial loss 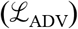. Lazarou^24^ combined both the generative and autoencoding aspects into an Autoencoding Generative Adversarial Network (AEGAN). The major limitation of AEGAN however, is that it requires two discriminators, an encoder, and a generator - so supporting large model architectures becomes difficult to implement in practice since the memory and compute required to train becomes technically infeasible and/or too expensive^25^.

An innovative way to take advantage of the ideal properties of both GANs and AEs is to combine their functionality. Pathak *et al*.,^26^ called this a Context Encoder (CE), whereby an image is first augmented to remove blocks of pixels, then processed through an AE, with 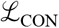 relative to the unmodified input image. This forces the model to create a semantically meaningful representation of the missing data (*i.e*. GAN) while learning a latent representation (*i.e*. AE). However, CEs have only been applied to smaller images (512×512)^27^ that are unrelated to the ultra-high resolution required for computational pathology.

Here, we combine CE with the progressive framework to encode high resolution images from digital pathology and bias this learning toward encoding only non-diseased skin tissue so that the model will not correctly encode abnormal tissue - forcing high reconstruction error that can be exploited for anomaly detection. We are able to show that our Progressive Context Encoders for Anomaly Detection (P-CEAD) model can segment tumor regions with high accuracy without the need for manual segmentation by a pathologist - representing a major step forward for more complex digital pathology workflows. This new method was able to achieve 89% pixel-level accuracy for anomalous regions of interest when compared to manually segmented melanomas.

## METHOD

The training dataset consists of a total of 200 slides from the Department of Laboratory Medicine and Pathology at Mayo Clinic. A senior dermatopathologist selected them from skin excisions based on the absence of inflammation, neoplastic process, and quality of glass slides. The testing dataset includes 9 skin slides with definitive invasive melanoma. All slides are anonymized and scanned at 40X with Aperio ScanScope V1. The invasive melanoma in the test dataset was annotated by a pathologist using QuPath 0.2.3^28^.

Due to the complexity of the task and allowing for modularity, P-CEAD involves several phases of processing and model training (**FIGURE 1)**. Briefly, in Phase 0, the preprocessing step identifies and removes image patches that are highly similar in image content. In Phase 1, the progressive autoencoder with inpainting is trained using 150 normal WSIs. During Phase 2, inference is run on 25 normal WSIs to calculate the normal error reference distribution. Next, in Phase 3, reconstruction error profiles from 25 additional normal WSIs were compared to the reference distribution to determine the value of 3 standard deviations from normal, which is used as a binary threshold to flag pixels as anomalies. Finally, inference uses a smoothed and filtered kernel density estimator (KDE) on the binary pixel flags in nine tumor-containing WSIs to determine an appropriate KDE threshold for defining an anomalous region.

**Figure 1.**
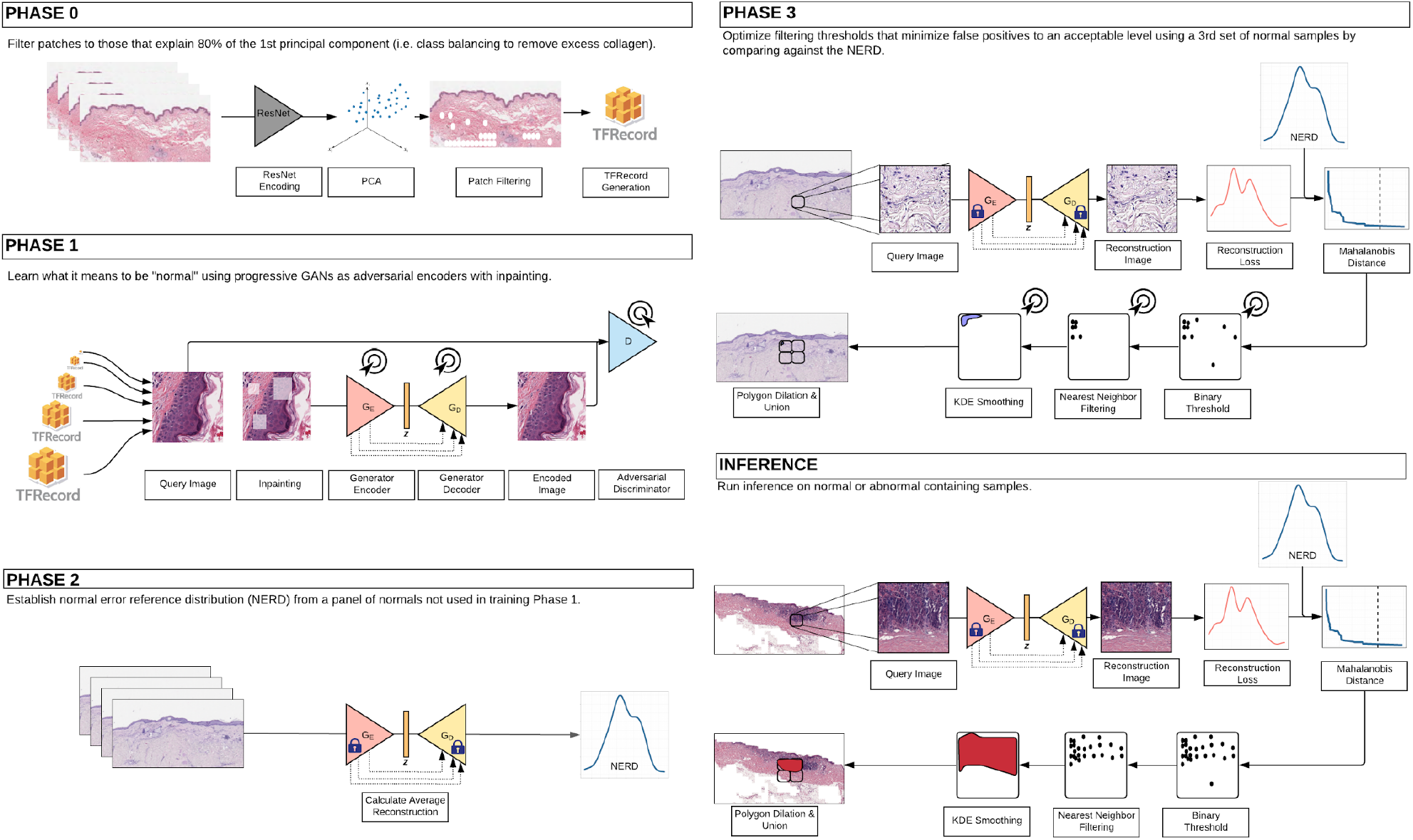
Overall architecture of the multiple components of P-CEAD.

### Phase 0: Preprocessing

The goal of the P-CEAD is to learn the manifold of normal images so it can identify outliers on that manifold for anomaly detection. Phases 1, 2, and 3 of P-CEAD all require normal images, so a significant amount of training data is necessary. However, many of the image patches contain redundant information. Skin sections are made of 3 compartments: epidermis, dermis, and subcutis. The subcutis is made of white fat appearing as mostly empty vacuoles in the H&E slide. The dermis is primarily made of collagen and represents an overwhelmingly large contribution of the total tissue observed on each slide. However, the epidermal layers (stratum corneum, lucidum, granulosum, spinosum, basale) represent the vast majority of diagnostically relevant regions. The class imbalance of these three layers means that the networks would mostly see collagen and fat rather than learning to focus on the epidermis. To overcome this limitation, training images are pre-filtered. First each image is passed through ResNet50 to obtain a vector representation of the image. Principal Component Analysis (PCA) is then applied to this collection of image vectors to project the vectors into an orthonormal basis. From there, the images corresponding to the first 80% of the first principal component were selected. As expected, manual inspection also confirmed that these selected patches captured mostly epidermis, borders, and examples from each of the layers, whereas many images with collagen and whitespace were removed. Across the 200 WSIs there were 534531 image patches in the pre-filtering set and 427625 post-filtering (80% of the original image patch count). However, the filtering was not uniformly 80% for each slide with some slides having very little filtering and others being extremely filtered. The minimum slide filtering had 99.29% of image patches remaining and the maximum slide filtering had 9.71% of image patches remaining, with an average of 76.1%.

Since P-CEAD involves three training phases, three distinct training sets were generated to ensure that the same image patches were not used for multiple phases. Training Phase 1 was the most computationally intensive, the majority of WSIs (n=150, 330415 patches) were used in this Phase. Twenty five WSIs each were used for training Phases 2 and 3, corresponding to 44693 and 52517 patches, respectively. As mentioned previously, these “normal” slides were selected based on the absence of inflammatory and neoplastic processes.

### Phase 1: Network weight training

Before training the network weights, images must be augmented. Otherwise, due to the architecture (and even more specifically the skip connections), the generator would simply become a compressive autoencoder. However, by masking the inputs the generator is forced to learn not only how to encode and decode the image, but also constrain the encoding and decoding to be context specific - in essence learning to encode images in a way that enforces an expected image type (in this case an normal skin WSI).

A percentage of pixels in each image (default 20%) are masked for each image patch. Random squares are generated using a halving geometric series such that the total sum of all of the masked pixels approximately matches the desired masking percentage. The squares are allowed to overlap, and if so, will randomly have their intersections unmasked, creating more complex shapes. This is uncommon enough that the effective mask percentage is still close to the setpoint, yet it does help break edge symmetries to discourage the model from just learning edge detection. Each mask block then randomly shuffles its pixels so that the original information is still within the mask region, albeit not in the correct spatial order. Randomly shuffled pixels produced more realistic inpainting than standard black pixel masking, but increased processing time when shuffling on-the-fly for each image patch. Adding randomly shuffled masked versions of the image patches to the TF Records would decrease the amount of compute and memory cost, but doing it on-the-fly ensures that each epoch will produce a different masking for each image which increases the diversity for learning the normal image manifold and reduces the likelihood of overfitting on a fixed set of masked images. Masked images are only used during the Phase 1 of training, whereas the other phases of training only use the unmasked image patches.

The model architecture at a high level consists of a generator and a discriminator. Having an additional network to act as an encoder (as in some GAN architectures) yielded negligible improvement and increased training time so was excluded from the final model architecture. All subnetworks follow the general pattern of Kerras *et al*.’s Progressively Growing GANs such as the generator using pixel normalization, the discriminator using minibatch standard deviation, WGAN loss with gradient penalty, epsilon drift penalty, no batch normalization, leaky ReLUs for activations, grows in resolution by factors of two from 4×4 images to 1024×1024 images, Adam optimizer with β_1_= 0 and β_2_= 0.99, etc.

The generator has a bowtie autoencoder structure consisting of an encoder that takes an image, x, as input and ZX outputs a latent vector, z, and a decoder which takes a latent vector, z, as input and outputs an image 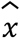. During Phase 1, the actual input to the bowtie is x’, the masked version of image x. To help facilitate high-resolution image generation, the bowtie architecture is specifically a U-net with skip connections between the encoder and decoder’s corresponding convolutional layers. Without skip connections, high resolution generated images were blurry from high resolution pixel information being lost at the U-net bottleneck. Skip connections were also pruned during testing, but much of the high resolution detail was lost by not including all of them and thus all skip connections are retained in the final model.

The discriminator is a standard progressively growing GAN discriminator. However, conditional weights were added for reconstruction and adversarial loss terms. For the generator, the final weighting scheme was *L_con* = *1.0 * L2(x - G(x))* and *L_adv* = *1.0 * -q(D(G(x)))* giving the total generator loss of *L_gen* = *L_con + L_adv = L2(x-G(x)) - ExpVal(D(G(x)))*. For the discriminator there were two adversarial loss terms *1.0 * D(x) - 1.0 * D(G(x)).* In addition there is a WGAN gradient penalty (GP) (γ)^29^ and an epsilon drift penalty (EDP) (ε) as done in Kerras *et al.* giving the total discriminator loss of *L_dis* = *q(D(x)) - q(D(G(x))) +* γ *+* ε.

For each step in training Phase 1, for each image *x* in a minibatch of image patches *X,* random masked blocks were created for each patch using the shuffling method. These augmented images *x’* are the same tensor shape as the original input images *x*. *x’* is then passed through the bowtie, with intermediate tensors flowing along the skip connections. The final output of the generator’s decoder will be an image 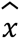, of the same tensor shape as the generator input *x’*, and therefore the same as our original inputs from the TF Records. The L2 norm between the original images *x* and the generated images 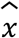 is the generator’s reconstruction loss term.

Both the real images *x* and the generated images 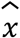 then proceed to the discriminator network where each example will receive one logit indicating whether the discriminator thinks it is the original image or is augmented. The discriminator will then be updated using the mean of the difference between the real and fake logits in addition to the GP and EDP. The negative mean of the fake logits is then added to the generator network using alternating gradient descent, since that provides more stable learning than simultaneous gradient descent. Both the discriminator and the generator had only one network weight update at each step.

Phase 1 was trained for 1 epochs at each image size. The prediction outputs after training the Phase 1 model are the generated images 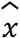 and the absolute errors between the input images *x* and the generated images 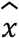.

### Phase 2: Error distribution calculation (NERD)

Once the progressive GAN’s network weights have been trained, the model weights are fixed in the bowtie generator. Beginning here, images are no longer augmented with masks since the inpainting training through the learned network weights is complete. The purpose of this phase is to learn the Normal Error Reference Distribution (NERD). This comes from the maximum likelihood estimation of the errors which are assumed to be gaussian in nature. Therefore the NERD is a multivariate gaussian distribution of the absolute errors between *x* and 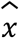 with a mean vector for each color channel (RGB) and a 3×3 covariance matrix for the color channels. The NERD’s mean vector and covariance matrix parameters are calculated from 25 WSIs not used in Phase 1.

The motivation for including the NERD is due to the fact that all imperfect models produce prediction errors and those errors are samples from a distribution of errors. Normal images should produce very low prediction errors of a certain distribution due to the progressive autoencoding in Phase 1, assuming the learned manifold of normal images is accurate. On the other hand, anomalous images should produce higher prediction errors, representing a different distribution.

The prediction outputs after training this phase of the model are Mahalanobis distances, calculated using the color channel means and covariance matrix. This distance metric is commonly used for finding multivariate outliers, whereas unlike Euclidean distance treats each axis independently (a sphere), Mahalanobis distance considers the scales (and cross correlations) of each axis (an ellipsoid). Lower distances correspond to pixels that have low absolute errors between input and generated images *x* and 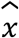, respectively, while larger distances correspond to pixels that have larger absolute errors.

### Phase 3: Dynamic distance threshold, KDE smoothing, and dilation

Knowing the Mahalanobis distances, one can determine whether an observed value exceeds that from the expected distribution. Larger distances are indicative of higher reconstruction error and are more likely to be anomalous. To convert the linear measurement of distance into a binary classification metric such as anomalous or not, a threshold can be determined where anything above it is assigned “anomalous” and everything below it is assigned “normal”. Using the fixed bowtie and the NERD, The optimal binary flag threshold is determined from a final catlog of 25 normal WSIs. Each image patch is run through the bowtie generator to yield the reconstruction error profile and compared with the NERD to calculate each pixel’s Mahalanobis distance. Just like for the NERD, these Mahalanobis distances form a distribution across all of the images in the Phase 3 training dataset, so the mean and standard deviation of Mahalanobis distances are calculated for this set. The threshold is set to be equal to the mean plus a number of standard deviations for the Phase 3 training set.

At full resolution, each image patch is a matrix of 1024×1024 pixels, with each pixel having a value of 0 (normal) or 1 (anomaly) which equates to over a million pixels. Up to this point, all pixels have been treated as independent. However, even at 40X magnification, each pixel represents a region smaller than the nucleus of a single cell - far below the resolution of the human eye, much less the resolution that a human would be capable of providing manual annotation for which to make benchmark comparisons. Moreover, there is nothing inherently intuitive or medically relevant for a single pixel. Instead, larger regions are of interest. Given these constraints, it is unreasonable to expect every pixel in a true anomaly to be flagged as an anomaly. Instead, spatial information can be leveraged to more accurately define the anomalous regions.

Since pixel-level flags can be noisy, spurious anomalous pixels in an otherwise large normal pixel region may incorrectly be classified as anomalies. To account for these spurious calls, two steps are performed: filtering and smoothing. To remove false positive pixels, we want to remove any small clusters of pixels. Looking at each flagged pixel, a cluster is removed if within a specified connectivity (default = 1) that the neighboring flagged pixel count is less than minimum adjacent pixel neighborhood size (default = 5). In other words, given an anomalous pixel *a*, if pixels connected to it and pixels connected to those and so on add up to less than 5 pixels surrounding *a*, then *A* is removed, otherwise it is retained. After regional filtering, a patch-level threshold is also applied, requiring that at a minimum of *A* (default = 10) anomalous pixels are found, otherwise all anomalous flagged pixels will be removed.

For the remaining pixel-level anomaly flagged images that need to be smoothed, a two dimensional gaussian kernel density estimator (KDE) is used on the pixel-level anomaly flags. Other kernel types and several bandwidth values were explored, but the best results were obtained with a gaussian kernel with a bandwidth of 100. A minmax normalization then transforms those evaluations to be within [0., 1.] and then scaled by *(anomaly_flag_counts /scaling_factor) ^ scaling_power* for visual consistency. These will be used for thresholding at a specified value between [0., 1.] to create a boolean mask. Therefore, rather than using the scaled kernel densities with a color map, we directly use those as is, making a grayscale KDE image.

Finally, the scaled grayscale KDE images can be compared against a specified threshold (default = 0.2) to create a bitmask image. These are then rasterized and converted to polygon objects using Shapely^30^ which are then dilated (default=4) to expand the shapely polygon beyond the image patch bounds. A final union of polygon objects is rendered across all image patches in a WSI. In this way, potential edge effects from neighboring patches will be eliminated. We then intersect the unioned polygons with a polygon of the original patch boundaries to constrain any dilated exteriors from extending outside the patch regions.

To measure performance relative to human annotation, the intersection, union, and difference between the two sets of polygons are used to create resultant polygon sets that represent different confusion metrics. True positives (TP) are measured as the area of intersection between the KDE and human-curated annotation polygons. False positives (FP) are measured as the area of difference between the KDE polygons and the true positive intersection polygons. False negatives (FN) on the other hand are measured as the area of difference between the annotation polygons and the true positive intersection polygons. This leaves the true negatives (TN) as the remaining area of the original patches. Sensitivity is calculated as TP / (TP + FN) and specificity as TN / (TN + FP).

## RESULTS

Phase 0 computation took 12.5 hours of computation to compress with ResNet and run PCA using 2GB of memory serially in a Google Colab notebook instance with 32 vCPUs and 208 GB RAM and 2 NVIDIA Tesla T4s. All distributed training was performed on n1-highmem-16 virtual machines with two NVIDIA Tesla V100 GPUS on the Google Cloud Platform (GCP). Phase 1 training required 129.5 hours using 16GB memory. Phase 2 and 3 both completed in under 1 hour with 16GB memory.

We initially attempted to simply adapt the GANanomalyDetection architecture to whole slide images. However, the approach failed to yield acceptable images and was discarded. When exploring images generated by the GAN, it was apparent that it had only learned to encode collagen and fat, which make up the vast majority of pixels in whole slide images of skin. We attempted to reduce the redundant images by selecting the images that most contributed (80%) to the overall variance using PCA (*see Methods*). However, we could not filter too many images out or else the model would not be exposed to sufficient examples of normal tissue. Another issue we found with GANanomalyDetection was that it either learned to encode the real images or the fake images - but never both. As a result, either the fake or the real images appeared blurry or would have a mode collapse when using the loss calculation *G(z)-(G(E(G(z))* (**Supplemental Figure 1**). This could not be overcome despite applying multiple weights to each loss function or changing combinations thereof.

As an alternative approach, we pivoted from a GAN-based architecture to an Adversarial Autoencoder-based architecture. Real images were encoded and decoded well in terms of histological structure, but distortions in color were also present (**Supplemental Figure 2**). This coloration artifact was overcome by adding skip connections to the autoencoder, producing a more reasonable reconstruction. The downside of the Autoencoder was that it simply learned to encode images - regardless of whether they contained anomalies, so input regions were masked from the image to learn context encoding. The rationale is that the context encoder would learn to preferentially compress and decompress images free of anomalies because it learned what normal should look like. When asked to compress and decompress a region with anomalies, the reconstruction error should be much higher in abnormal images because the model would not have learned how to encode and decode abnormal images (**FIGURE 2**).

**Figure 2.**
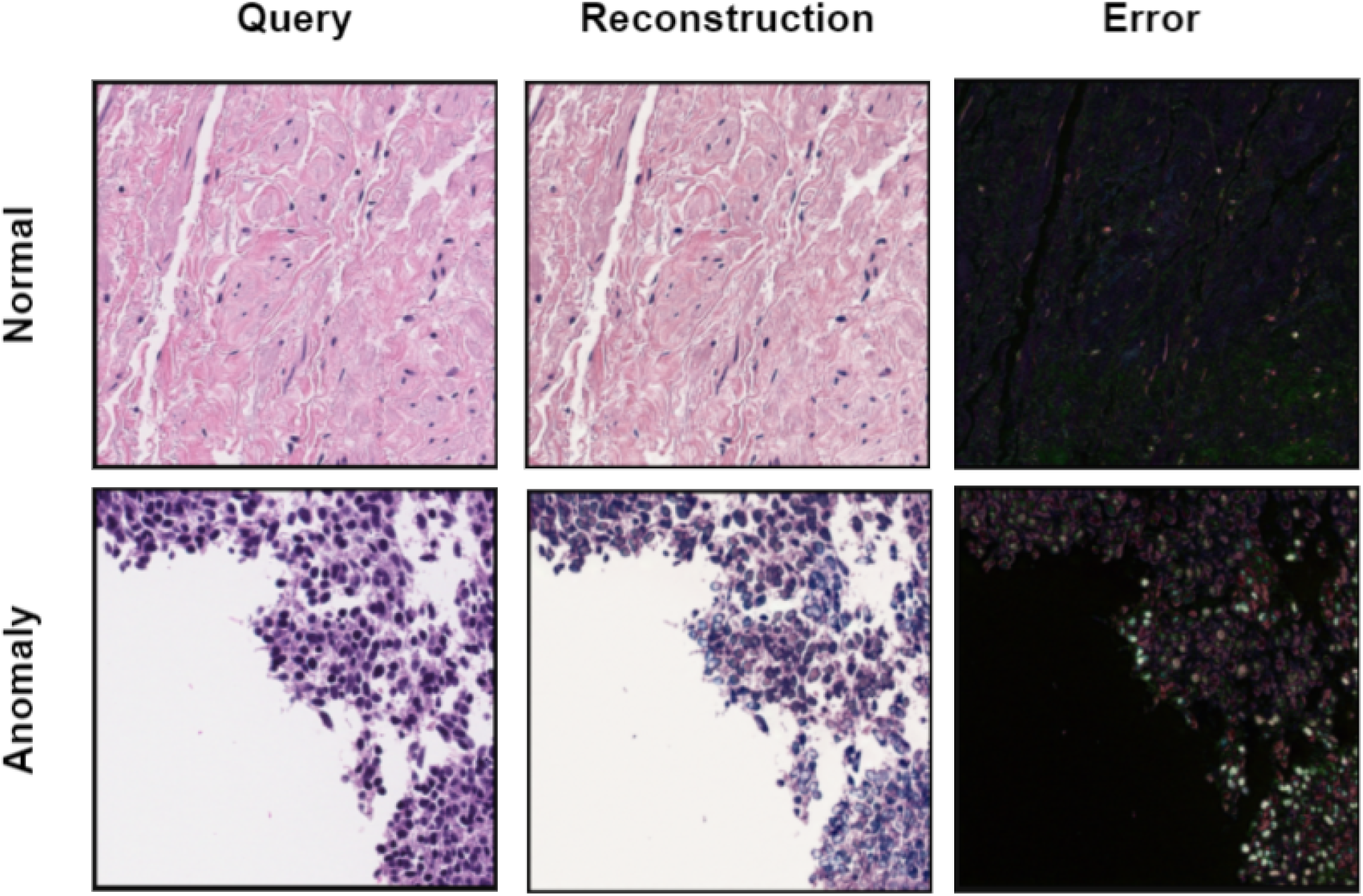
Examples of reconstruction error from image reconstruction.

Once satisfied with the overall approach, we evaluated model performance on 9 slides with anomalies that were not in any phase of training. A board certified pathologist created ground truth segmentation maps to indicate areas of melanoma on H&E slides. Using a KDE threshold of 0.21 and a polygon dilation factor of 4, on average the model was both sensitive (94% ± 8), specific (87% ± 7), and accurate (89% ± 7)(**TABLE 1**).

**Table 1.**
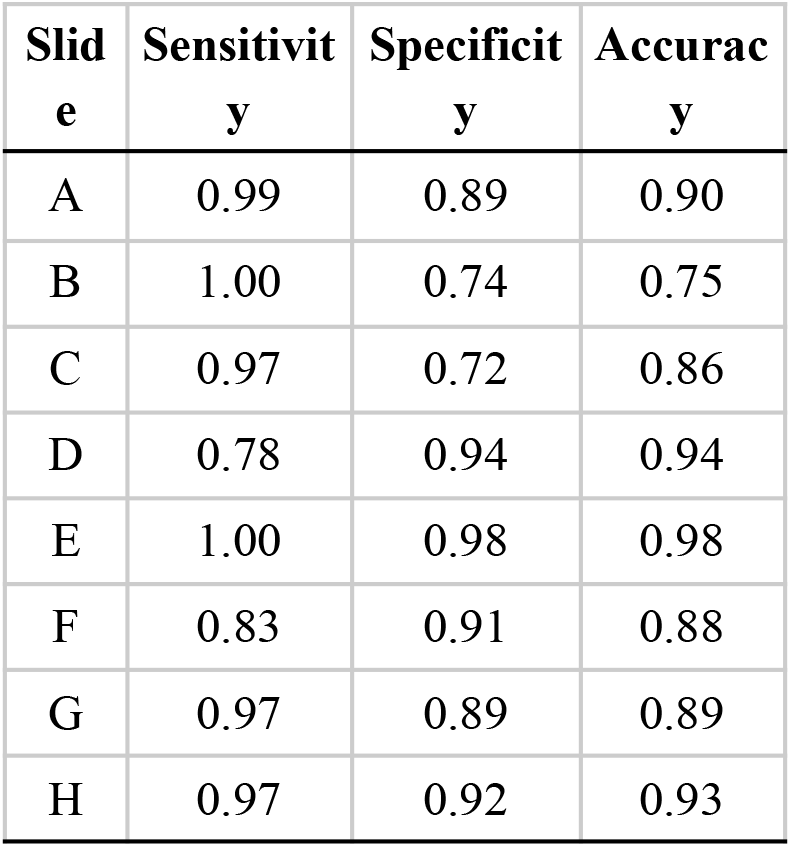
Performance metrics for 9 whole slide images containing melanoma.

| Slide | Sensitivity | Specificity | Accuracy |
| --- | --- | --- | --- |
| A | 0.99 | 0.89 | 0.90 |
| B | 1.00 | 0.74 | 0.75 |
| C | 0.97 | 0.72 | 0.86 |
| D | 0.78 | 0.94 | 0.94 |
| E | 1.00 | 0.98 | 0.98 |
| F | 0.83 | 0.91 | 0.88 |
| G | 0.97 | 0.89 | 0.89 |
| H | 0.97 | 0.92 | 0.93 |

We should also note that inference parameters for identifying melanoma may not be the same for non-melanoma lesions or anomalies in other tissues. Rather, the inference module we constructed allows users to select, adapt, and change filtering criteria to suit the task at hand through configurations. Each modification to the inference filters would necessarily alter the performance metrics. For example, choosing an alternative dilation factor for the polygon expansion can have a dramatic effect on sensitivity and specificity (**FIGURE 3**).

**Figure 3.**
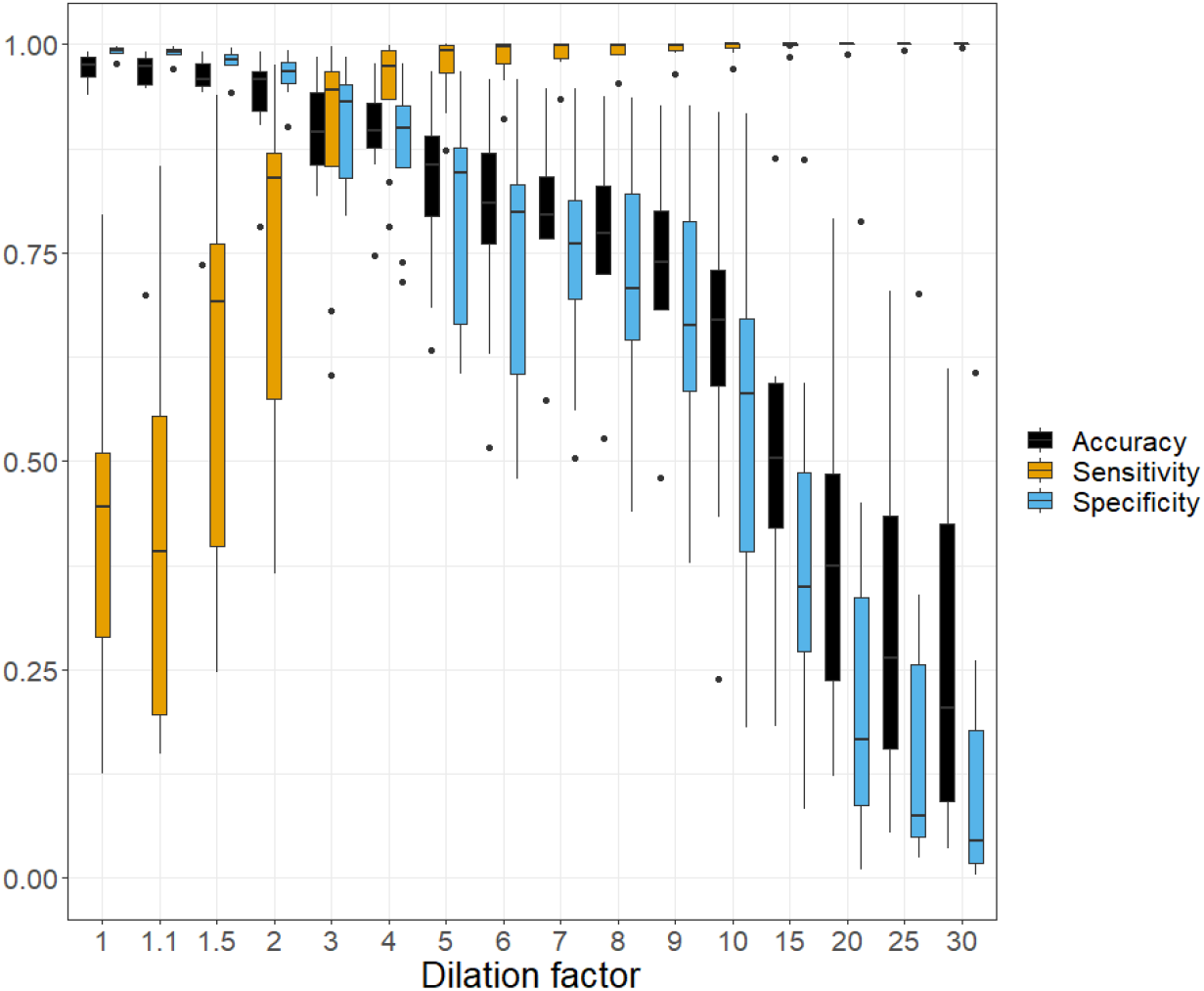
Performance metrics for 9 whole slide images as a function of polygon dilation.

### Exploration of predictions

To improve our understanding of the P-CEAD model, we explored predictions at the patch level. **FIGURE 4** shows representative images that contain true and false positives and negatives. The top row is an example of normal whereas the bottom is completely involved with melanoma. In the normal image, most of the predictions were correctly identified as not containing anomalies, but pixels toward the top of the image patch were. Upon closer inspection, this is thought to be the result of epidermal lymphocytic infiltrate or presence of keratinocytes with abundant clear cytoplasm. Such changes, while not associated with tumors, were limited in our training set of histologically normal samples, for which very few were expected to have extensive immune infiltrate. Review of additional false positive regions suggested an enrichment in areas where the model expected - but did not identify - a preserved epidermis, due to the presence of parakeratosis, intraepidermal lymphocytes, knife artefact, epidermal denudation, or lumina of large arteries.

**Figure 4.**
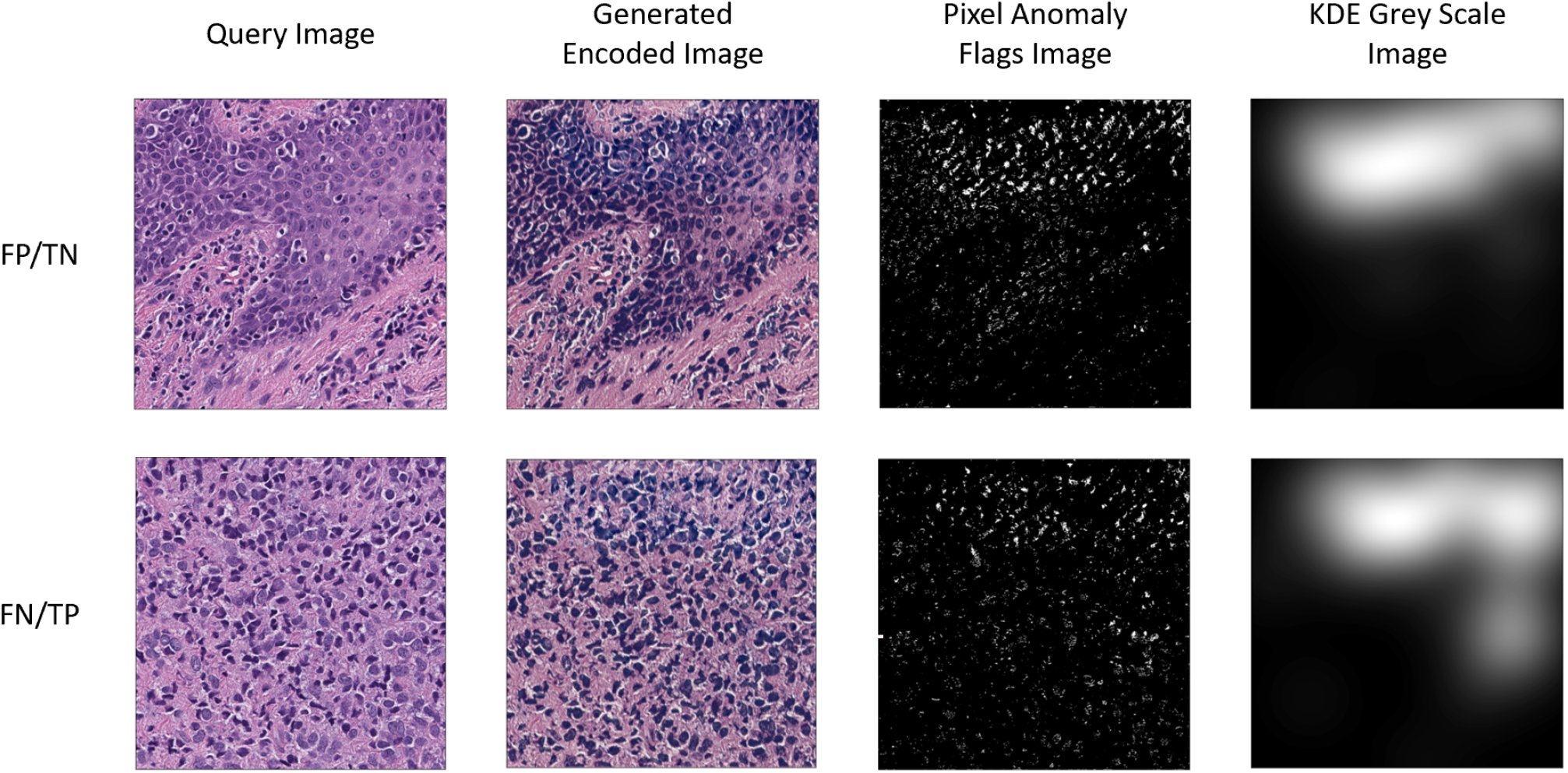
Examples of correct and incorrect predictions from P-CEAD. (Top) The query image contains only normal tissue. Reconstructing the image through the P-CEAD model results in an overall darker image, which also corresponds to a higher error rate and subsequent flagging of individual pixels. The greyscale image defines the region for polygon creation, with white being used to call anomalies. (Bottom) This query image contains 100% tumor, but only a portion was flagged as anomalous. This type of model exploration can inform users of how and where filters could be applied to refine final predictions. FP, false positive; TN, true negative; FN, false negative; TP, true positive.

The case of false negatives is shown in the bottom row of **FIGURE 4**. Here, the entire image is involved with melanoma, yet only a small portion of the image patch (Upper right corner) is flagged after filtering. Our interpretation of the missed pixels is due to an increase of degenerated tumoral cells, as well as increased extracellular matrix. Like the other example, reconstructed images are generally darker in color than the original query image - which ultimately results in higher reconstruction errors and flagged as anomalies.

## DISCUSSION/CONCLUSION

The main contributions to P-CEAD are the diversity sampling for unsupervised patch selection, addition of inpainting, removal of unnecessary loss terms from previous architectures, and the development of a modular secondary process for fine tuning anomaly detection.

Image inpainting was required for *G_E_* to preferentially encode patterns that were observed only in the training set. RGB values from blocks of randomly chosen coordinates were removed or shuffled before being autoencoded. The reconstruction loss (measuring the per-pixel delta of the autoencoded image relative to the pre-augmented image) was thus not only measuring the quality of compression/decompression, but also the ability of *G_E_* to “hallucinate” realistic patterns of normal. Inpainting was imperative to bias the AE such that the AE would produce larger errors in the reconstructed images in the area of patterns it had not observed before (i.e. anomalies).

Multiple architectures and combinations of weighted loss terms were attempted before developing P-CEAD. Models, including GANanomalyDetection, were able to encode real or generated images, but never both. We attributed this effect to the latent-space loss term 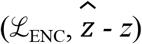 and its dependency on a high quality generator architecture. Otherwise, as in the case of GANanomalyDetection, the training step is trying to minimize the loss when *z’* is dependent on the outputs of three independent models (generator-encoder, generator-decoder, and an additional encoder). In P-CEAD, the 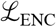 is removed and thus obviates the need for the additional encoder network and now more resembles an autoencoder for the generator of the GAN system. However, unlike autoencoders (adversarial or otherwise), we are not interested in the latent space representation^31^. Instead, the generative component occurs though inpainting during *G_E_* and *G_D_* using the 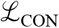 for optimization. The addition of skip-connections also had a profound influence on the reconstructed image quality and added to the model’s generative ability.

P-CEAD defines anomalies in a more innovative and practical way than previous methods. GANomaly^18^, f-AnoGAN^32^, EBGAN^16^, and GANanomalyDetection^20^ all define their anomaly scores as some derivation of reconstruction error relative to the query image. In contrast, P-CEAD measures the difference in reconstruction error relative to the NERD. This distinction is subtle, but important. In our approach, we separate the model training from error calculation. In an ideal world, one would have captured all representations of normal images during model training - but this is rarely possible in reality. The model we have presented here was trained using SVS files from an Aperio scanner with a JPEG2000 image compression that was sectioned, stained, and imaged at Mayo Clinic. We cannot be certain that the reconstruction error distribution from a slide processed by an external lab on a different scanner would be the same as ours. However, having established the NERD on the internal dataset, one can easily compare it to the reconstruction error from the external laboratory’s normal slides. If the distributions are the same, then no assumptions are violated and the model should behave as expected. If they are not the same, then normal samples from the external lab could be used to calculate a new NERD for processing the external data, without retraining the computationally demanding Phases 0 and 1.

A key limitation to P-CEAD is that it does not define what anomalies are. This is a separate task that should be performed after P-CEAD has segmented any anomalous regions. Another limitation is the nearest neighbor filtering during inference. In theory, requiring a minimum number of pixels to be co-located will decrease the analytical sensitivity of the method. Also, the costs and technical considerations required for Phase 1 training may be prohibitive for many clinical departments.

Despite its limitations, P-CEAD represents a significant step forward for applying artificial intelligence to whole slide images within a clinically relevant context. Unlike challenge competitions or toy problems, P-CEAD is not based on a simple classification of tumor vs normal. Many clinically benign nevi mimic the architecture of pathogenic varieties - which could otherwise be classified as tumor in a binary prediction algorithm. P-CEAD allows the “grey area” to exist without overconfident claims of diagnostic accuracy. Furthermore, P-CEAD may have additional uses such as labeling anomalous regions for tissue scraping and downstream molecular testing.

## ACKNOWLEDGEMENTS

This work was funded by the Leon Lowenstein Foundation (SNH), the Mayo Clinic/Google Joint Steering Committee, and the Mayo Clinic Center for Digital Health (SNH, TF).

## AUTHOR CONTRIBUTIONS

RG, QG, SP, NP developed and tested the code, documentation, and experiments. CM and TF performed clinical annotation and case selection. SNH and TF designed the study, oversaw the study execution, and secured its funding. All authors discussed the results and commented on the manuscript and approved the final version.

## SUPPLEMENTAL METHODS

**Table S1.**
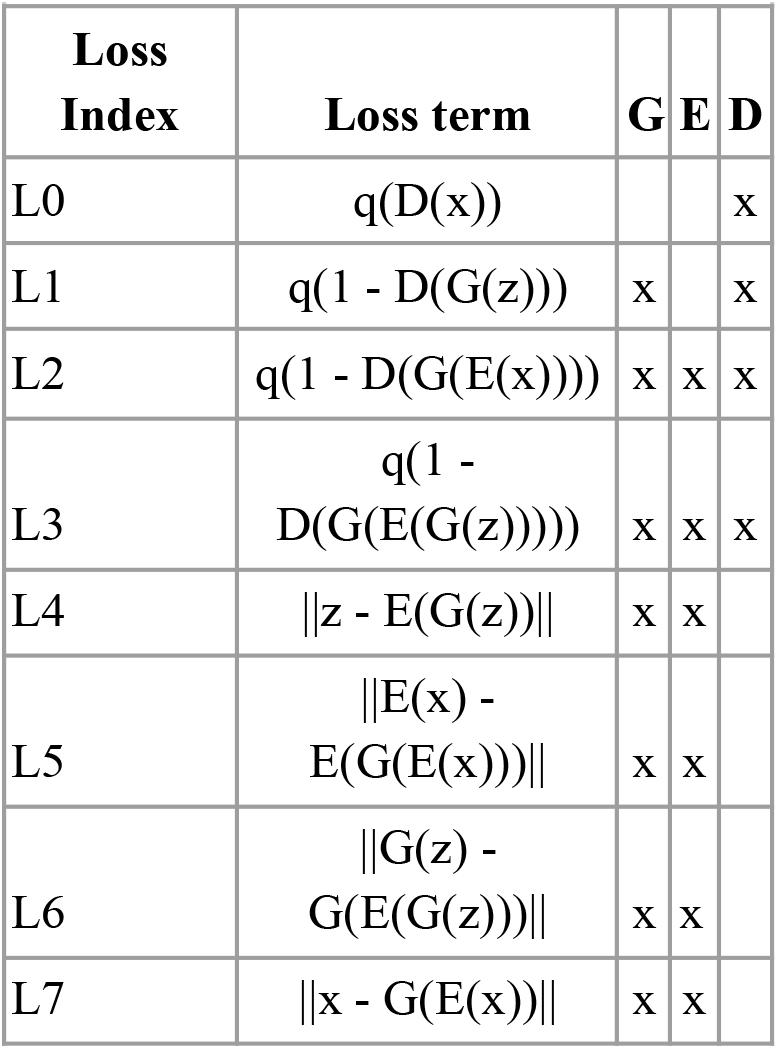
Loss tem combinations.

### Loss Functions

When exploring the triple network architectures (as in GANanomalyDetection) eight different weighted losses were attempted (Table S1). The use of those losses could be applied to any or all of the independent networks and have a weight of 1, 0.01, 0.001, or 50. In general, we observed when L6 and L7 were greater than 0.001, then the generated images were visually similar to normal skin. Manual review of the generated images showed dominant representation of fat and collagen, despite being selected against in pre-training. L2 and L3 had little influence when applied to the encoder or discriminator architecture with respect to generating more realistic images. The real limitation was when trying to encode real images using the encoder network - particularly in images containing epidermal layers. Real images in this context were consistently highly blurred, rendering them useless for further investigation - despite a stabilization of all the loss terms.

**Supplemental Figure 1.**
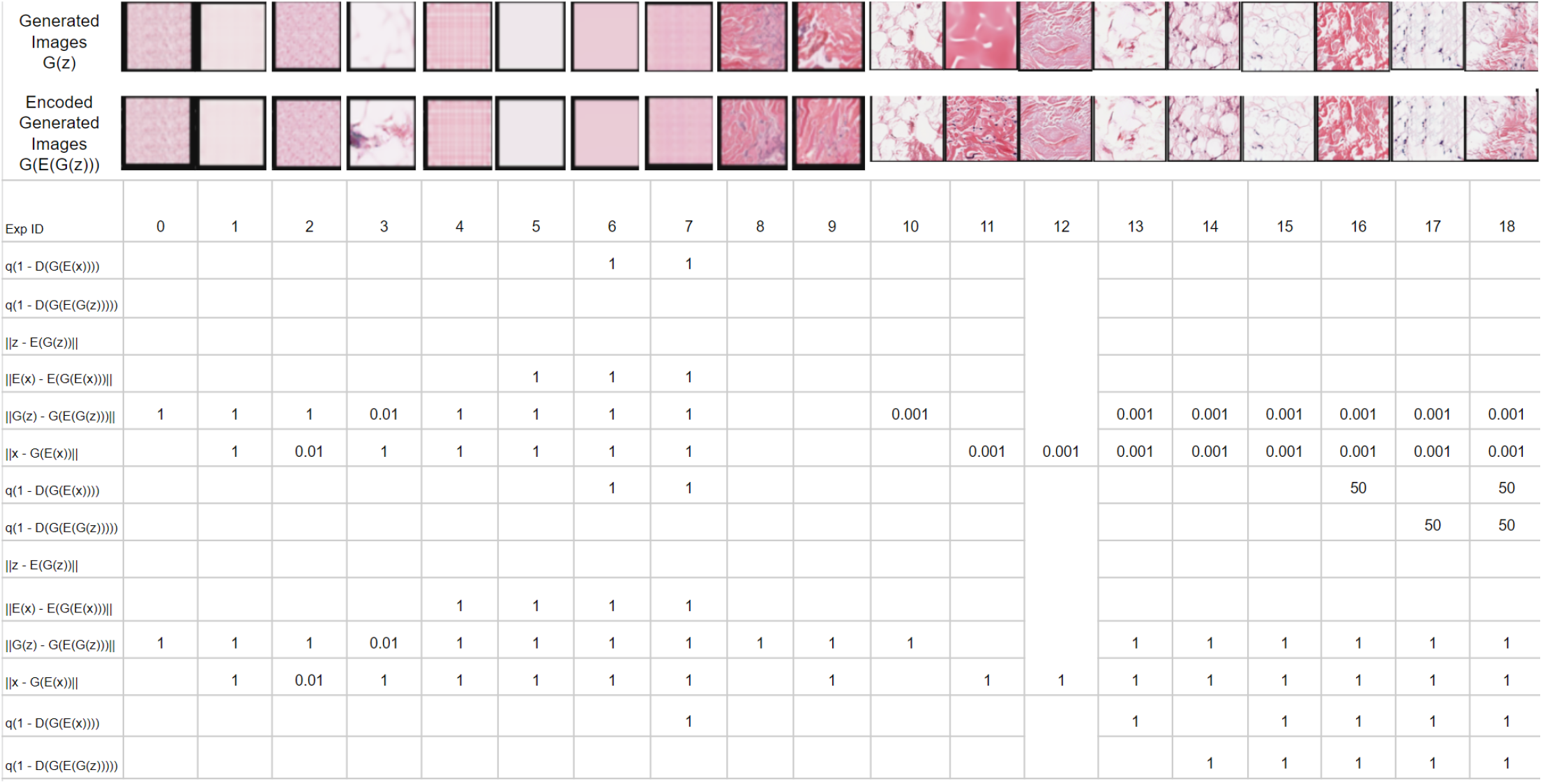
Loss function penalty exploration

### Caveats

#### Inappropriate image inputs

If the network weights are trained on images that contain anomalies, then since the training is unsupervised the model has no ability to know that these images aren’t normal and therefore the learned manifold expands to encode those images within it. Likewise, the error distribution training in phase 2 needs to contain only normal type images. This time, if anomalous images are within the training dataset, since the manifold is now fixed due to the first phase of training being complete, it is the error distribution’s parameters that will now expand. This is due to the off-manifold anomalous images generating larger errors which then get encoded into the error distribution. Lastly, phase 3’s training requires all images to be normal so that the tightest distance threshold can be set. If anomalous images were accidentally included in its training dataset, then larger distances would result, ergo raising the acceptable distance threshold. Thus, since all three training require the training datasets to have as little anomalous example contamination, it is crucial to ensure that any images that could be contaminated are filtered out before the training phases.

#### Different PCA thresholds

Other global filtering percentages were also assessed. Exceeding 80% resulted in many blurry and blank image patches being included in the training datasets. Being stricter than 80% didn’t inherently make the images unsuitable for training, but it did throw out too many useful images which could lead to our model not having enough unique data to learn what “normal” looks like across the three training phases. Thus we stuck with keeping the top 80% of image patches.

#### Mahalanobis Distance thresholding

One could directly use the Mahalanobis distance as a numerical score for pseudo likelihood of being normal. Distances are in the range of 0-Inf with lower scores meaning more likely normal. However, each pixel’s distance is independent of its neighbors - even though there should be a very strong spatial correlation. This is the logical basis for the KDE smoothing since we would not expect pixels near each other to be representing different anomalies. However, the kernel density estimator expects a two-dimensional array binary mask, so distances do not satisfy this condition. This results in the kernel density estimator scoring each sample the same and therefore since there is no variance every sample results in a zero-valued grayscale pixel, thus the resulting image is entirely zeros. This can be remedied by converting the input images into binary masks via a threshold, clipping, etc. which is what we do with the dynamic threshold learned in training phase 3.

**Supplementary Figure 2.**
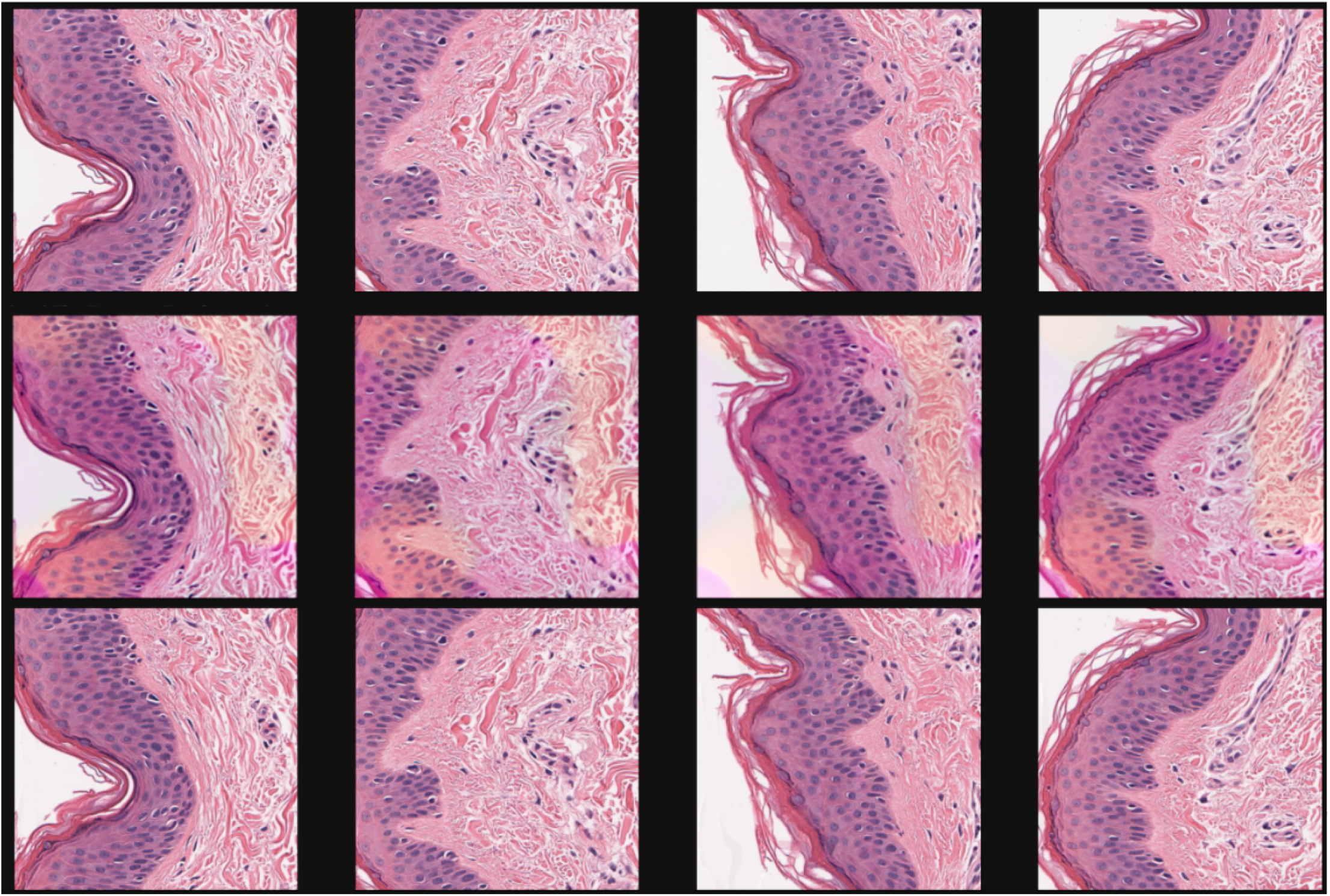
Effect of Skip Connections. (Top Row) Original image patch, (Middle Row) Skip connections turned off, (Bottom Row) Skip connections turned on. No uniform noise was added to *z* or to fake images, and the loss for ‘x_minus_G_of_x_l2_loss_weight’ was zero.

